# A cell-free transcription-translation pipeline for recreating methylation patterns boosts DNA transformation in bacteria

**DOI:** 10.1101/2023.09.16.557782

**Authors:** Justin M. Vento, Deniz Durmusoglu, Tianyu Li, Sean Sullivan, Fani Ttofali, John van Schaik, Yanying Yu, Lars Barquist, Nathan Crook, Chase L. Beisel

**Affiliations:** Department of Chemical and Biomolecular Engineering North Carolina State University, Raleigh, NC 27695, USA; Helmholtz Institute for RNA-based Infection Research (HIRI), Helmholtz Center for Infection Research (HZI), 97080 Würzburg, Germany; Medical Faculty University of Würzburg, 97080 Würzburg, Germany

**Keywords:** bacteria / cell-free / methylation / restriction / transformation / TXTL

## Abstract

The bacterial world offers diverse strains for understanding medical and environmental processes and for engineering synthetic-biology chasses. However, genetically manipulating these strains has faced a long-standing bottleneck: how to efficiently transform DNA. Here we report IMPRINT, a generalized, rapid and scalable approach based on cell-free transcription-translation (TXTL) systems to overcome DNA restriction, a prominent barrier to transformation. IMPRINT utilizes TXTL to express DNA methyltransferases from the bacterial host’s restriction-modification systems. The expressed methyltransferases then methylate DNA *in vitro* to match the host DNA’s methylation pattern, circumventing restriction and enhancing transformation. With IMPRINT, we efficiently multiplex methylation by diverse DNA methyltransferases and enhance plasmid transformation in gram-negative and gram-positive bacteria. We also developed a high-throughput pipeline that identifies the most consequential methyltransferases, and we apply IMPRINT to facilitate a library screen for translational rules in a hard-to-transform Bifidobacterium. Overall, IMPRINT can enhance DNA transformation, enabling use of increasingly sophisticated genetic manipulation tools across the bacterial world.

## INTRODUCTION

While physical barriers such as the cell wall can interfere with transformation, arguably the most common barrier is posed by restriction-modification (R-M) systems^1^. These systems are one of a growing set of defenses^2^ but represent the most prevalent across bacteria. To confer immunity, R-M systems rely on DNA methyltransferases (MTases) that methylate genomic DNA with a unique methylation pattern using S-adenosyl methionine (SAM) as a methyl donor as well as restriction endonucleases (REases) that cleave DNA with foreign patterns. Four types have been defined based on the need for a specificity protein to guide the MTase and REase (Type I), the MTase and REase acting independently (Type II), the MTase and REase forming a complex capable of both methylation and restriction (Type III), and the REase cleaving methylated DNA (Type IV)^3^. Transformed DNA possessing the wrong methylation pattern undergoes extensive cleavage, resulting in a massive drop in the number of transformed colonies. As a further challenge, bacteria often possess not one but multiple R-M systems conferring unique methylation patterns that can vary even between related strains^4,5^.

Recreating these patterns or mutating the short recognition sites have proven to be an effective way to restore transformation, in some cases radically boosting transformation^5–8^. However, existing approaches have been heavily constrained in their ability to fully reproduce these patterns in a rapid and comprehensive means (**Table S1**). For example, approaches involving *in vitro* methylation of DNA with purified MTases or by expressing the MTases in a plasmid-propagating strain of *E. coli* are laborious and suffer from methylation-induced cytotoxicity^5,8,9^, typically limiting their use to one or two DNA MTases that often provide incomplete protection^5,10^. Lysates of host cells can also be used to methylate DNA with the appropriate pattern, although methylation is highly inefficient and can lead to DNA degradation by host endonucleases^11–14^. Finally, an approach to identify and mutate MTase sites requires methylome sequencing followed by resynthesizing entire DNA constructs, where many of these mutations could interfere with plasmid maintenance or the expression and function of the encoded constructs^7^. Therefore, new approaches are needed to simplify how R-M systems are circumvented across bacteria.

## DESIGN

To tackle this challenge, we sought to harness TXTL as a distinct means to overcome DNA restriction. TXTL recapitulates transcription and translation in a lysate or solution of purified components, allowing the functional expression of any RNA and protein in minutes to hours without the need for cell culturing or protein purification^15^. We envisioned a TXTL-based pipeline in which DNA MTases identified in a host bacterium (e.g. through bioinformatics or listed on the REBASE database^16^) are expressed and combined with DNA in a single methylation reaction, and the resulting DNA is purified and transformed into the host. DNA lacking any methylation (e.g. PCR product, plasmid DNA extracted from a MTase-free strain) serves as the starting point to immediately circumvent Type IV R-M systems that cleave methylated DNA^8^. We call the resulting pipeline IMPRINT, for Imitating Methylation Patterns Rapidly IN TXTL (**Fig. 1a**). With IMPRINT, methylation by multiple MTases, including Type I MTases that also require expressing a specificity protein, is straightforward and completed in a day. Cytotoxicity concerns are also minimized, as replicating cells are not required for plasmid methylation, and linear chemically-synthesized DNA can be used to express the MTases.

**Fig. 1:**
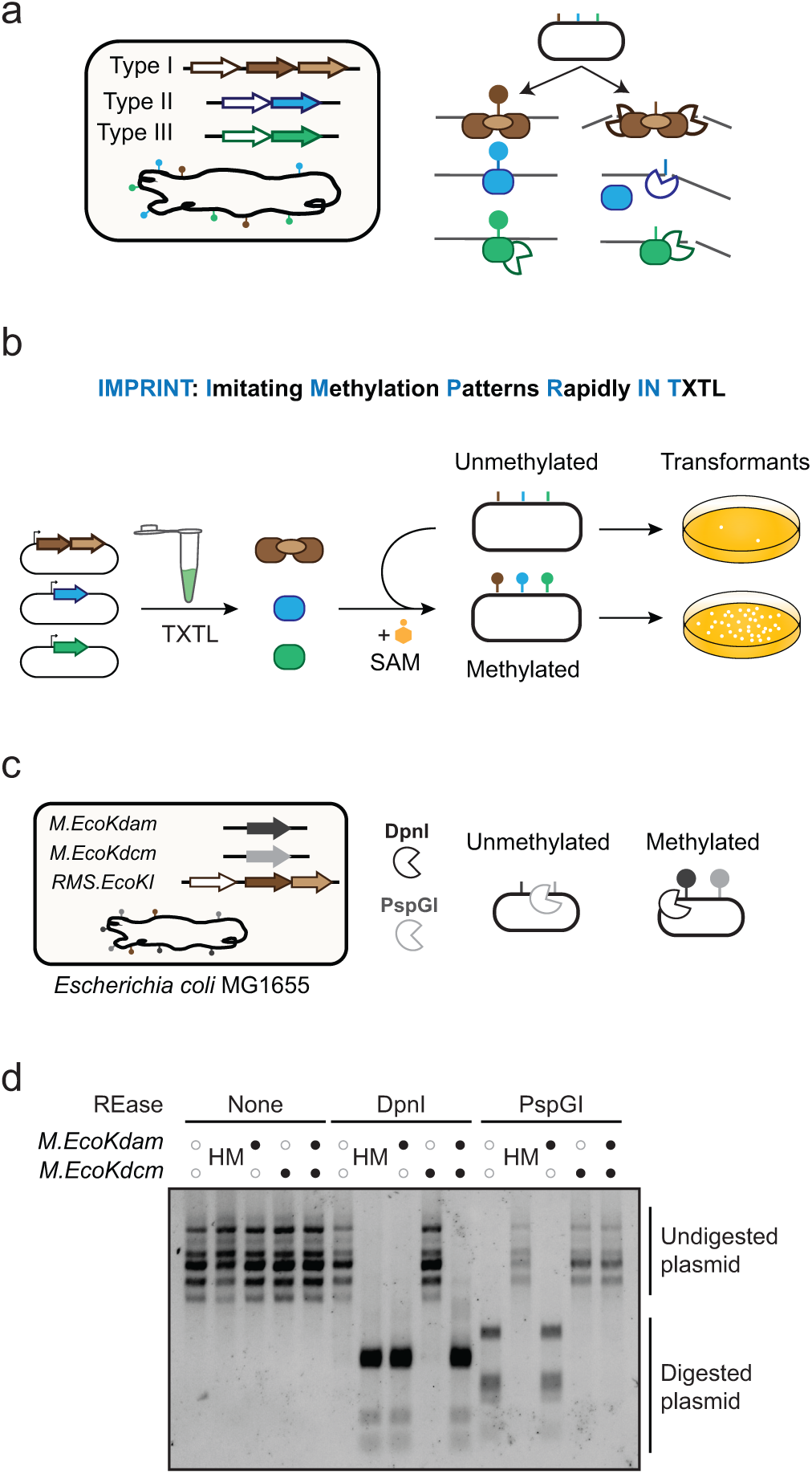
**A TXTL-based pipeline imbues shuttle vectors with a host’s methylation pattern to overcome restriction barriers in bacteria**. **a,** Overview of the different types of R-M systems that cleave unmethylated DNA. MTases (darker shaded arrows) methylate DNA, while REases cleave unmethylated DNA (white arrows). Type I systems rely on a specificity protein (lighter shaded arrow) to bind the DNA recognition site. **b,** The cell-free pipeline to recreate host methylation patterns. MTases from the host strain are cloned into expression plasmids and individually expressed in TXTL. The expressed MTases are then combined to methylate initially unmethylated DNA that can then be purified and transformed into the host strain. **c,** Assessing TXTL-based expression of Dam (encoded by *M.EcoKdam*) and Dcm (encoded by *M.EcoKdcm*) from *E. coli* MG1655. The REase DpnI cleaves sites methylated by Dam, while the REase PspGI cleaves sites not methylated by Dcm. **d,** Restriction digestion of plasmid pJV24 methylated by Dam and/or Dcm using IMPRINT. HM: host-methylated plasmid DNA extracted from *E. coli* MG1655 and subjected to IMPRINT without any added MTases.

## RESULTS

### Efficient methylation of plasmid and linear DNA with the *E. coli* MTases Dam and Dcm expressed in TXTL

We initially prototyped IMPRINT by expressing MTases from *E. coli*. Standard laboratory strains of *E. coli* (e.g. MG1655) possess two orphan DNA MTases, Dam (M.EcoKdam) and Dcm (M.EcoKdcm), which respectively methylate adenine in GATC and the second cytosine in CC(A/T)GG. To assess methylation, we used the restriction enzyme DpnI, which cuts GATC when methylated by Dam, and PspGI, which cuts CC(A/T)GG unless it is methylated by Dcm (**Fig. 1b**). Each MTase was cloned in an *E. coli* strain with an inhibited P70a promoter to avoid expression of the potentially cytotoxic MTase. After expressing MTases in a TXTL reaction, the reaction was then incubated individually or together with the *E. coli* plasmid pJV24 containing 10 Dam recognition sites and 7 Dcm recognition sites. After completing IMPRINT, DpnI digested the plasmid incubated with Dam or both Dam and Dcm, while PspGI was unable to digest the plasmid incubated with Dcm or both Dam and Dcm (**Fig. 1c**). We obtained similar digestion patterns with the plasmid extracted from *E. coli* MG1655, which expresses both MTases. Together, complete digestion with DpnI and complete protection from PspGI affirmed efficient methylation with both MTases. Efficient methylation was further achieved with Dam encoded on a linear DNA construct or when methylating linear DNA (**Fig. S1**), establishing that linear constructs can be used to express MTases and to undergo methylation through IMPRINT. These results show that IMPRINT can be used to methylate different forms of DNA with MTases expressed from linear or plasmid constructs in TXTL.

### Applying IMPRINT to host MTases in *Salmonella* enhances plasmid transformation

The next major question was whether DNA methylated through IMPRINT could enhance transformation. We turned to the LT2 strain of the gram-negative pathogen *Salmonella enterica*, which is related to *E. coli* but restricts *E. coli* DNA through its set of well-characterized R-M systems^17^. As listed on REBASE, the associated MTases include two orphan MTases Dam (M.SenLT2dam) and Dcm (M.SenLT2dcm) homologous to those in *E. coli*, another orphan MTase (M.SenLT2IV), a Type I MTase requiring a specificity protein (MS.SenLT2II), and a Type III MTase (M.SenLT2I) (**Fig. 2a**)^18^. The five MTases were cloned into expression plasmids, with the MTase and specificity genes for the Type I R-M system cloned as an operon. We further selected two transformation plasmids (the 2.5-kb JV400 and the 14-kb JV414) harboring a different number of recognition sites for each MTase and different modes of replication (**Fig. 2b**). For the three MTases with compatible REases (M.SenLT2dam with DpnI, M.SenLT2dcm with PspGI, and M.SenLT2IV with NsiI), we confirmed methylation by each MTase based on altered cleavage patterns by the corresponding REase (**Fig. S2A-C**). Methylation by the Type III MTase was confirmed by cloning a REase site at the methylation site (**Fig. S2D**). We then electroporated either pJV400 or pJV414 methylated with different MTase combinations into LT2 (**Fig. 2c**). Compared to the unmethylated plasmid, none of the orphan MTases significantly enhanced transformation. In contrast, the Type I MTase and specificity protein provided a large increase in transformation (131-fold for JV400, 6-fold for JV414), while the Type III MTase provided a smaller but significant boost (6-fold for JV400, 1.5-fold for JV414). Importantly, combining both MTases in the same methylation reaction resulted in a much larger increase (1,000-fold for JV400, 32-fold for JV414) that approached the transformation levels for plasmid DNA extracted from LT2 possessing the strain’s complete methylation pattern (4,300-fold for JV400, 86-fold for JV414). Therefore, IMPRINT can employ multiple MTases from various R-M systems to enhance DNA transformation.

**Fig. 2:**
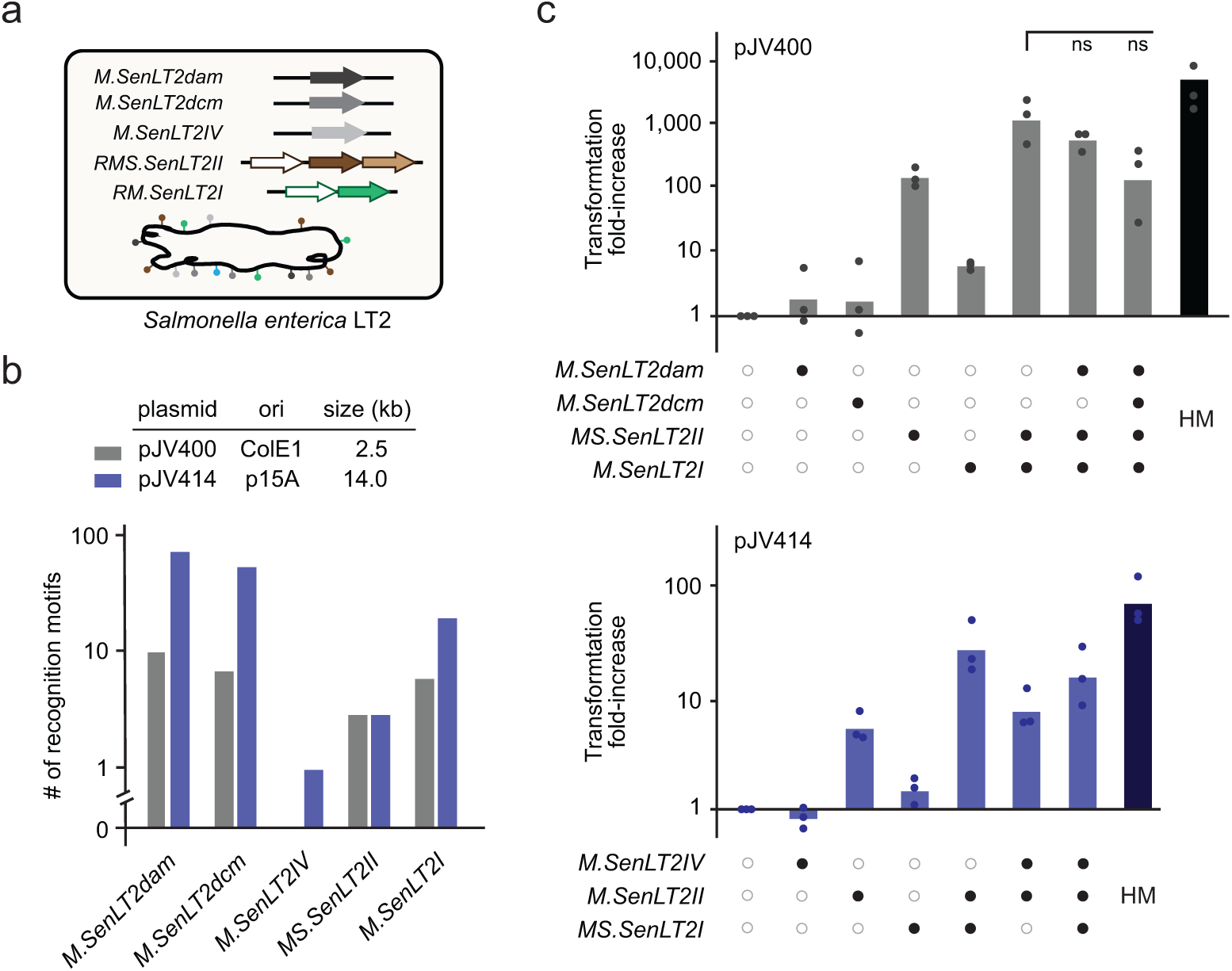
IMPRINT enhances plasmid transformation in the DNA-restricting *S. enterica* strain LT2. a, R-M systems associated with *S. enterica* LT2. MTases in gray are considered orphan MTases that are not associated with REases. **b,** Number of MTase sites present in the two tested plasmids. **c,** Electroporation efficiency of plasmids pJV400 (top) and pJV414 (bottom) subjected to IMPRINT with different MTase combinations into *S. enterica* LT2. All transformation efficiencies are normalized to that of unmethylated DNA. Error bars represent the geometric mean and standard deviation of triplicate independent experiments starting from separate TXTL reactions. The bottom of the bar marks the reference for statistical analysis. **: p < 0.01. *: p < 0.05. ns: p > 0.05. HM: host-methylated plasmid DNA extracted from *E. coli* MG1655 and subjected to IMPRINT without any added MTases.

### Removing the hemi-methylation dependence of the *Salmonella* Type I MTase enhances plasmid methylation and transformation

We noticed that including any of the orphan MTases with the Types I and III MTases reduced transformation (**Fig. 2c**). As this reduction could pose a limitation for bacteria harboring large numbers of R-M systems, we asked why transformation was reduced and if it could be restored. We first found that methylating plasmid DNA in a series of methylation reactions rather than a one-pot reaction restored transformation, suggesting that one of the MTases exhibited reduced activity (**Fig. S3a**) in the presence of the other MTases. When evaluating the rate of methylation with the Type I and III MTases, we noticed that the Type I MTase required a longer reaction time to achieve complete methylation based on transformation into LT2 (**Fig. S3b**). Similarly, the Type I MTase provided incomplete protection against cleavage of an introduced overlapping REase site, particularly in the presence of other MTases (**Fig. S3c**). One explanation for incomplete protection is that this subtype of MTases (Type IA) prefers hemimethylated substrates^19^, slowing methylation and allowing the other MTases to compete for available SAM. Fortuitously, prior work isolated mutants of the Type I MTase in *E. coli* (M.EcoKI) that readily accepted unmethylated substrates^20^ (**Fig. 3a**). Introducing two of these mutations into the Type I MTase from LT2 (**Fig. 3b**), we found that either mutation improved protection against HinFI cleavage from partial to complete even when including a second MTase (**Fig. 3c**), indicating more robust methylation of unmethylated DNA by the mutant MTases. The mutations also enhanced transformation into LT2 that was maintained when simultaneously methylating with Dam, Dcm, and the Type III MTase from LT2 (**Fig. 3d**) and allowed for shorter reaction times (**Fig. S4**). These results show that the Type I MTases can be a bottleneck in DNA methylation, although mutating the MTase to remove the preference for hemimethylated substrates can enhance transformation and allow the inclusion of other MTases in the methylation reaction.

**Fig. 3:**
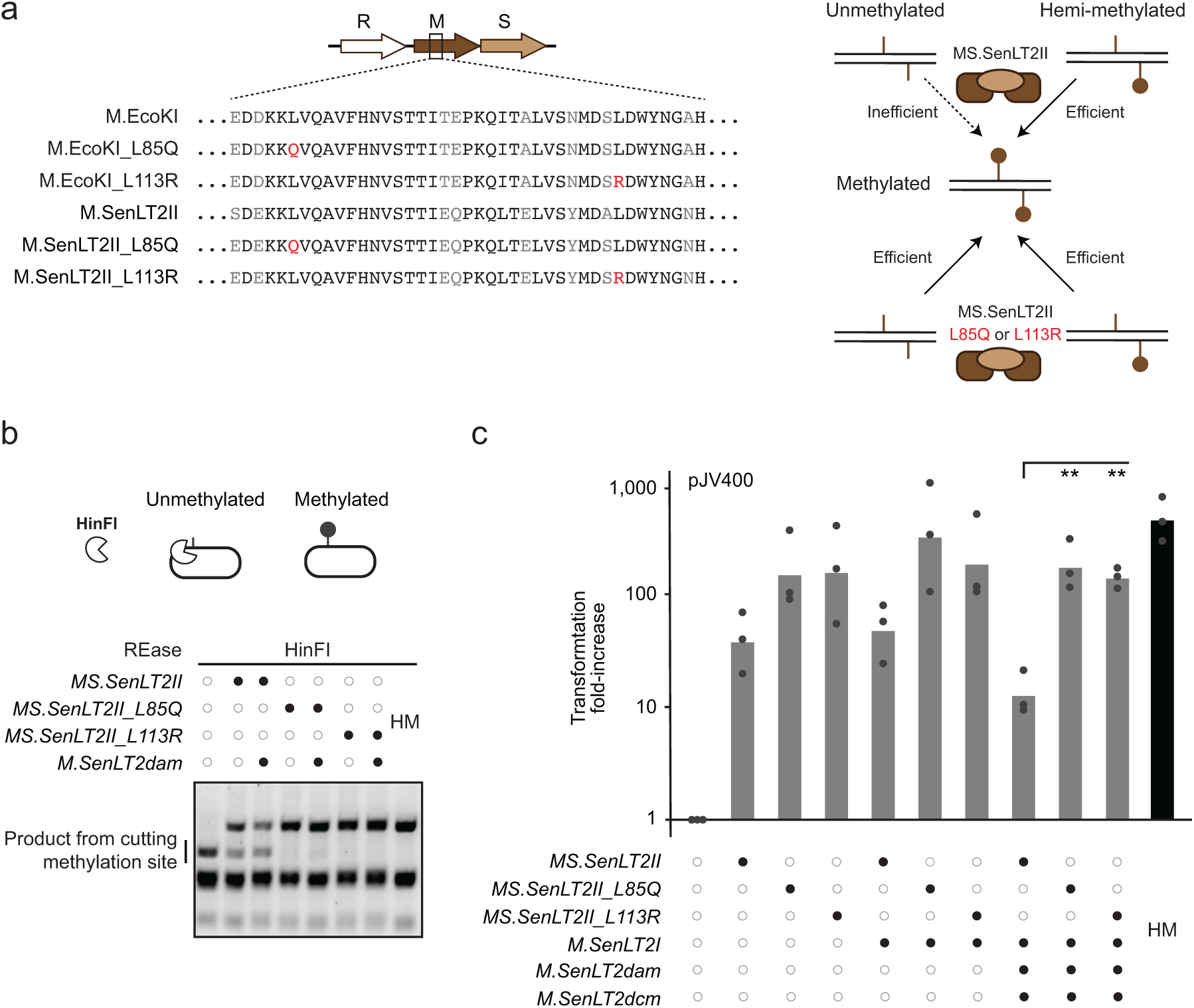
Conserved Type IA R-M methyltransferase mutations promote methylation of unmethylated DNA to achieve near-complete protection in *S. enterica* LT2. a, Aligned type IA MTases between *E. coli* and S. *enterica*. Mutations previously shown to enable efficient methylation of unmethylated DNA are shown in red. IA MTases normally prefer hemimethylated DNA as substrates. **b,** Restriction digestion of plasmid pJV412 methylated using IMPRINT. A HinFI restriction site was inserted to be methylated through one of the MS.SenLT2II recognition sites. HinFI digests the plasmid in six other locations, explaining the consistent lower bands on the gel. **c,** Electroporation efficiency of plasmid pJV400 subjected to IMPRINT with different MTase combinations into *S. enterica* LT2. All transformation efficiencies are normalized to that of unmethylated DNA. Error bars represent the geometric mean and standard deviation of triplicate independent experiments starting from separate TXTL reactions. The bottom of the bar marks the reference for statistical analysis. **: p < 0.01. *: p < 0.05. ns: p > 0.05. HM: host-methylated plasmid DNA extracted from *E. coli* MG1655 and subjected to IMPRINT without any added MTases.

### HT-IMPRINT allows high-throughput testing of MTase combinations

By applying different MTase combinations using IMPRINT, we determined that the Type I and III MTases contributed to enhanced DNA transformation in *S. enterica* LT2, while the orphan MTases played no role in enhanced transformation. For bacteria with a large number of R-M systems or much more laborious and intensive transformation procedures, testing the MTases individually or in combinations would be a long and involved process. We therefore sought to create a high-throughput version of IMPRINT that tests different combinations of MTases in a single transformation (**Fig. 4a**) to find the minimal set required for high transformation efficiencies. To reach this goal, we devised a setup in which a set of short barcodes are introduced into the shuttle plasmid. Several barcodes are then associated with one combination of MTases in case a given barcode introduces or removes a methylation site. The full set of barcodes representing all tested MTase combinations are then transformed into the host bacterium, and the transformed cells are plated or back-diluted into liquid medium. After isolating plasmid DNA from pooled colonies or liquid culture, the relative frequency of each barcode can be quantified by next-generation amplicon sequencing. The extent of barcode enrichment compared to the untransformed library would represent the boost in transformation provided by the associated MTase combination. We call this approach high-throughput IMPRINT (HT-IMPRINT).

**Fig. 4:**
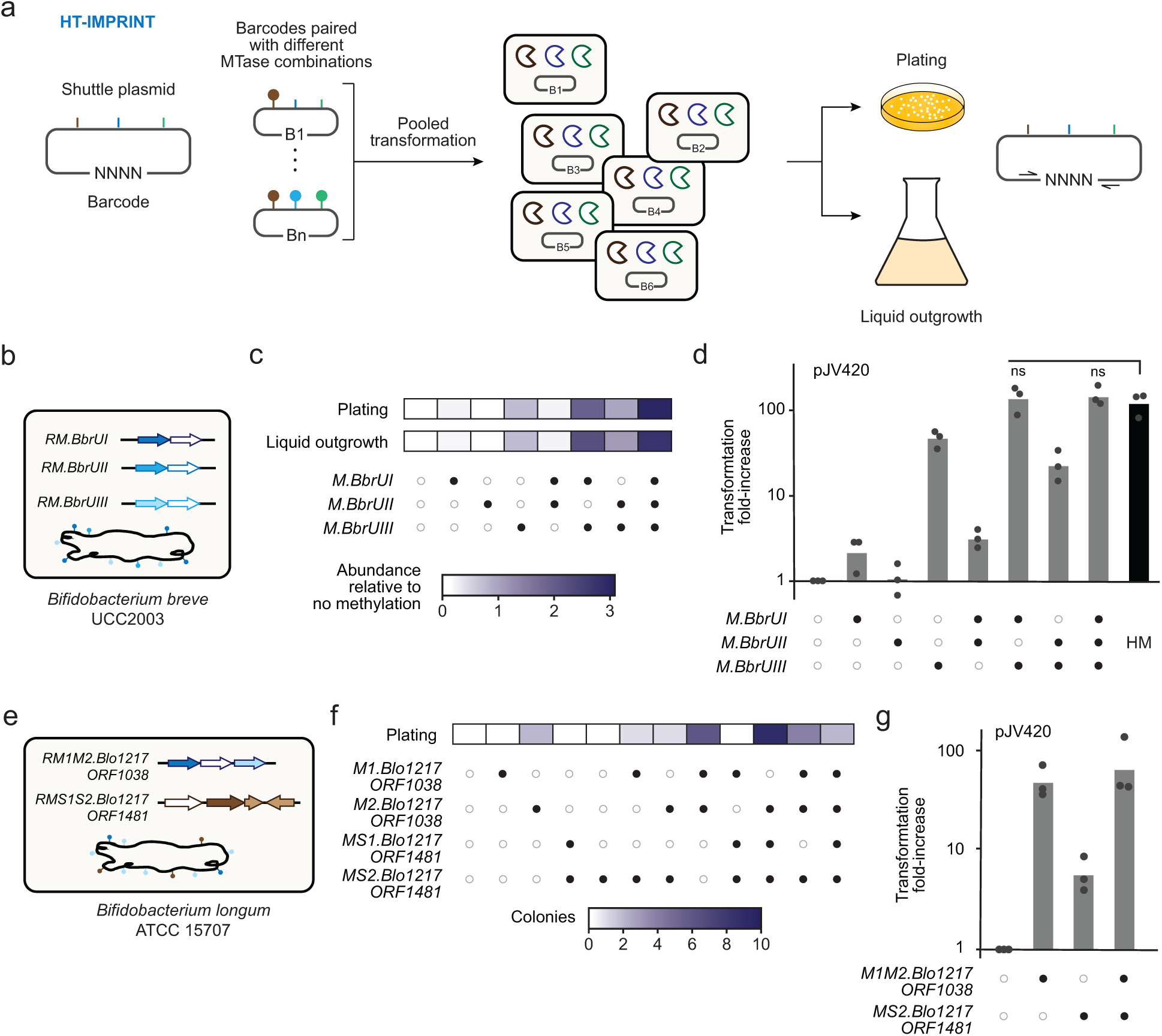
High-throughput IMPRINT determines optimal methylation patterns in recalcitrant Bifidobacteria. a, Overview of High-throughput IMPRINT (HT-IMPRINT). Barcodes are associated with each MTase combination to track how that combination impacts the transformation efficiency. **b,** R-M systems associated with *B. breve* UCC2003. **c,** Heat map of the relative impact of different MTase combinations using HT-IMPRINT with pJV420 in *B. breve* UCC2003. Three barcodes were associated with each MTase combination, where the reported values are the median relative abundance normalized to the untransformed library. **d,** Electroporation efficiency of plasmid pJV420 subjected to IMPRINT with different MTase combinations into *B. breve* UCC2003. **e,** R-M systems associated with *B. longum* ATCC 15707. **f,** Heat map of the relative impact of different MTase combinations using HT-IMPRINT with pJV420 in *B. longum* ATCC 15707. Two barcodes were associated with each MTase combination, where the reported values are the number of colonies possessing a barcode associated with the indicated MTase combination. **g,** Electroporation efficiency of plasmid pJV420 subjected to IMPRINT with different MTase combinations into *B. longum* ATCC 15707.

### HT-IMPRINT identifies MTase combinations that enhance plasmid transformation in different Bifidobacterial strains

To apply HT-IMPRINT and move beyond bacteria related to *E. coli*, we focused on Bifidobacteria. These gram-positive bacteria are common constituents of the human digestive tract, and many species are used as probiotics^21,22^. These bacteria are also strict anaerobes with more involved transformation procedures, possess a wide variety of R-M systems, and have proven difficult to transform^4,21^. We began with one strain, *Bifidobacterium breve* UCC2003, which harbors three characterized Type II R-M systems listed on REBASE and previously shown to interfere with DNA transformation (**Fig. 4b**)^23^. We cloned the three associated MTases (M.BbrUI, M.BbrUII, M.BbrUIII) into expression plasmids for IMPRINT, which were confirmed to methylate DNA in TXTL based on blocked cleavage of associated REases (**Fig. S5**). We then introduced a total of 24 unique barcodes into the *Bifiobacterium-E. coli* shuttle plasmid JV420 to cover the eight possible combinations of MTases (**Table S2**). Following IMPRINT with each MTase combination, the pooled plasmids were transformed into *B. breve* UC2003, and transformed DNA was recovered from pooled colonies or liquid culture followed by amplicon sequencing. The output from both approaches revealed that M.BbrUI and M.BbrUIII individually boosted transformation, with M.BbrUIII having a much stronger effect. Furthermore, combining all three MTases provided the greatest increase in transformation (**Fig. 4c**). Notably, there was very little variation among the three barcodes from each sample as a result of the screen (**Fig. S6).** Testing individual MTase combinations with IMPRINT yielded the same trend, where combining M.BbrUI and M.BbrUIII provided an increase in transformation (128-fold) similar to combining all three MTases (140-fold) (**Fig. 4d**). Importantly, the extent of transformation matched that for the shuttle plasmid extracted from *B. breve* UCC2003 (**Fig. 4d**), showing that DNA restriction was fully circumvented using IMPRINT. The results also matched the previously published role of M.BbrUIII in DNA restriction^23^, although our results show that including other MTases leads to an additive boost in transformation.

As a more challenging demonstration of HT-IMPRINT, we applied the technique to *Bifidobacterium longum* ATCC 15707. This strain contains two R-M systems listed on REBASE, where the Type II system encodes two different MTases (M_1_.Blo1217ORF1038, M_2_.Blo1217ORF1038) and the Type I system encodes an MTase with two different specificity proteins (MS_1_S_2_.Blo1217ORF1481) (**Fig. 4e**). This strain has also proven extremely difficult to transform^24^, and the R-M systems remain uncharacterized. Because of the lack of characterization, we cloned each MTase or MTase/specificity protein pair and immediately proceeded to HT-IMPRINT using the pJV420 shuttle plasmid, selecting 12 MTase combinations covered with 24 barcodes (**Table S3**). The resulting transformed cells did not grow in liquid culture once antibiotics were added, while we obtained 25 colonies following plating--both outcomes likely due to sub-optimal selection conditions (**Fig. 4f**). The colonies were associated with a mix of MTase combinations, where the most common MTases were M_1_.Blo1217ORF1038, M_2_.Blo1217ORF1038, and/or MS_2_.Blo1217ORF1481. Testing M_1_M_2_.Blo1217ORF1038 and

MS_2_.Blo1217ORF1481 individually and together using IMPRINT supported the output from HT-IMPRINT, where the greatest increase in transformation compared to unmethylated DNA came from the paired combination (74-fold) followed by M_1_M_2_.Blo1217ORF1038 (49-fold) and then MS_2_.Blo1217ORF1481 (6-fold) (**Fig. 4g**). The boost in transformation could not be compared to host-methylated DNA, as negligible DNA could be extracted from this strain. Overall, HT-IMPRINT allowed us to determine an effective combination of MTases for a recalcitrant strain of Bifidobacteria.

### Optimal MTase combinations are strain-specific and can reduce transformation in non-cognate strains

HT-IMPRINT helped reveal the most consequential MTase combinations for different Bifidobacterial strains. Ideally, the methylation pattern for one strain would improve transformation in other strains, easing the process of transforming diverse bacteria. However, these comparisons remain to be made in a systematic way. We therefore asked how the best MTase combination identified via IMPRINT in one strain impacts transformation in other strains. We selected the best minimal MTase combinations for *S. enterica* LT2, *B. breve* UCC2003, and *B. longum* ATCC 15707 and applied these to the *Bifidobacterium*-*E. coli* shuttle plasmid JV420 that can be propagated in all strains (**Fig. 5a**). Transforming this plasmid with each methylation pattern into each strain, we found that transformation was highest when the MTase combination was matched with the originating host strain (**Fig. 5b**). Interestingly, unmethylated DNA transformed more efficiently than DNA methylated with a pattern not from the host strain. The largest drop occurred using the *B. breve* methylation pattern to transform *B. longum*, showing that the methylation pattern of related species can be detrimental. Taking these data together, the best methylation pattern mapped to the originating strain, while transformation could be hindered by using an incorrect methylation pattern. The species-specific methylation pattern needed to enhance DNA transformation highlights the need for approaches tailored to individual bacteria, as afforded by IMPRINT.

**Fig. 5:**
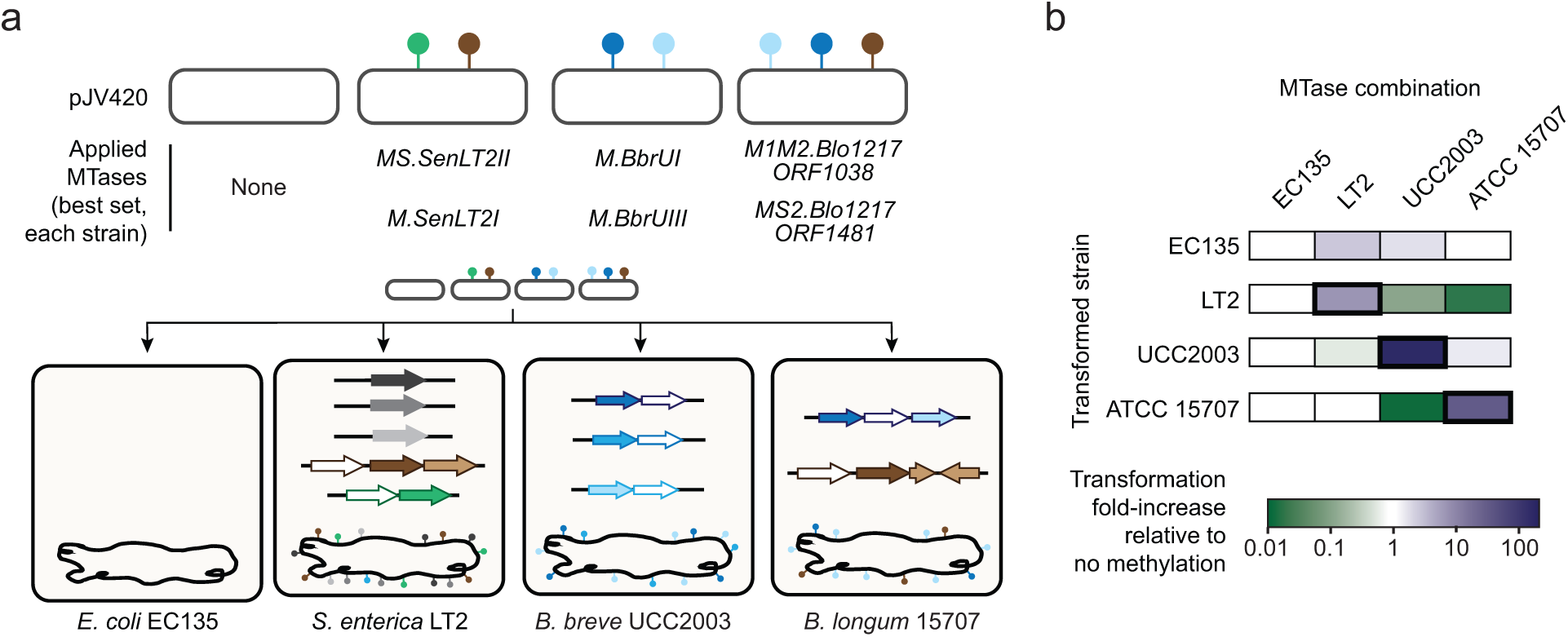
Identified MTase combinations that enhance plasmid transformation are strain-specific. a, Compatibility between optimal MTase combinations and transformation efficiency across bacterial strains. The plasmid pJV420 was methylated with optimal patterns for each strain using IMPRINT and transformed into the different strains. *E. coli* EC135 lacks any R-M systems and thus serves as a transformation control. **b,** Electroporation efficiency of pJV420 with the different methylation patterns into the different bacterial strains. All transformation efficiencies for a given strain are normalized to that of unmethylated DNA. Heat maps represent the geometric mean of triplicate transformations. See Fig. S7 for individual measurements. Error bars in d and g represent the geometric mean and standard deviation of triplicate independent experiments starting from separate TXTL reactions. The bottom of the bar marks the reference for statistical analysis. **: p < 0.01. *: p < 0.05. ns: p > 0.05.

### IMPRINT facilitates library-based interrogation of translational rules in *B. breve*

As a final demonstration of IMPRINT, we considered a general scenario in which maximizing the transformation efficiency is crucial: screening of large DNA libraries. Whether performing genetic screens, evolving proteins and pathways, or screening randomized DNA sequences for new functions, such procedures require obtaining not one transformant but enough transformants to encompass the entire library. Failing to do so limits the screening power of the procedure and confounds any downstream computational analyses.

To demonstrate how IMPRINT can enable screening of large DNA libraries in non-model bacteria, we devised a screen to elucidate rules governing protein translation strength in *B. breve* UCC2003. Within the *Bifidobacterium*-*E. coli* shuttle plasmid JV420, we randomized nine nts spanning the putative ribosome-binding site (RBS) and first nucleotide of the start codon of the tetracycline resistance gene *tetW* used to select for transformants in *B. breve* (**Fig. 6a**). Therefore, only those of the 262,144 unique sequences that yield sufficient expression of *tetW* to confer resistance to tetracycline would appear in the pool of transformants. A version of JV420 in which we deleted the RBS and start codon was used as the parent plasmid, which did not yield transformants of *B. breve*. Transforming the unmethylated library resulted in 8,200 transformants, or 3.1% coverage of the theoretical library (**Fig. 6b**). However, transforming the library methylated with M.BbrUI and M.BbrUIII resulted in an average of 28-fold increase in transformants, or 59.8% library coverage, over three independent experiments. We also isolated DNA from a liquid culture of the transformants without plating. This coverage offered a reasonable starting point to interrogate the resulting sequences.

**Fig. 6:**
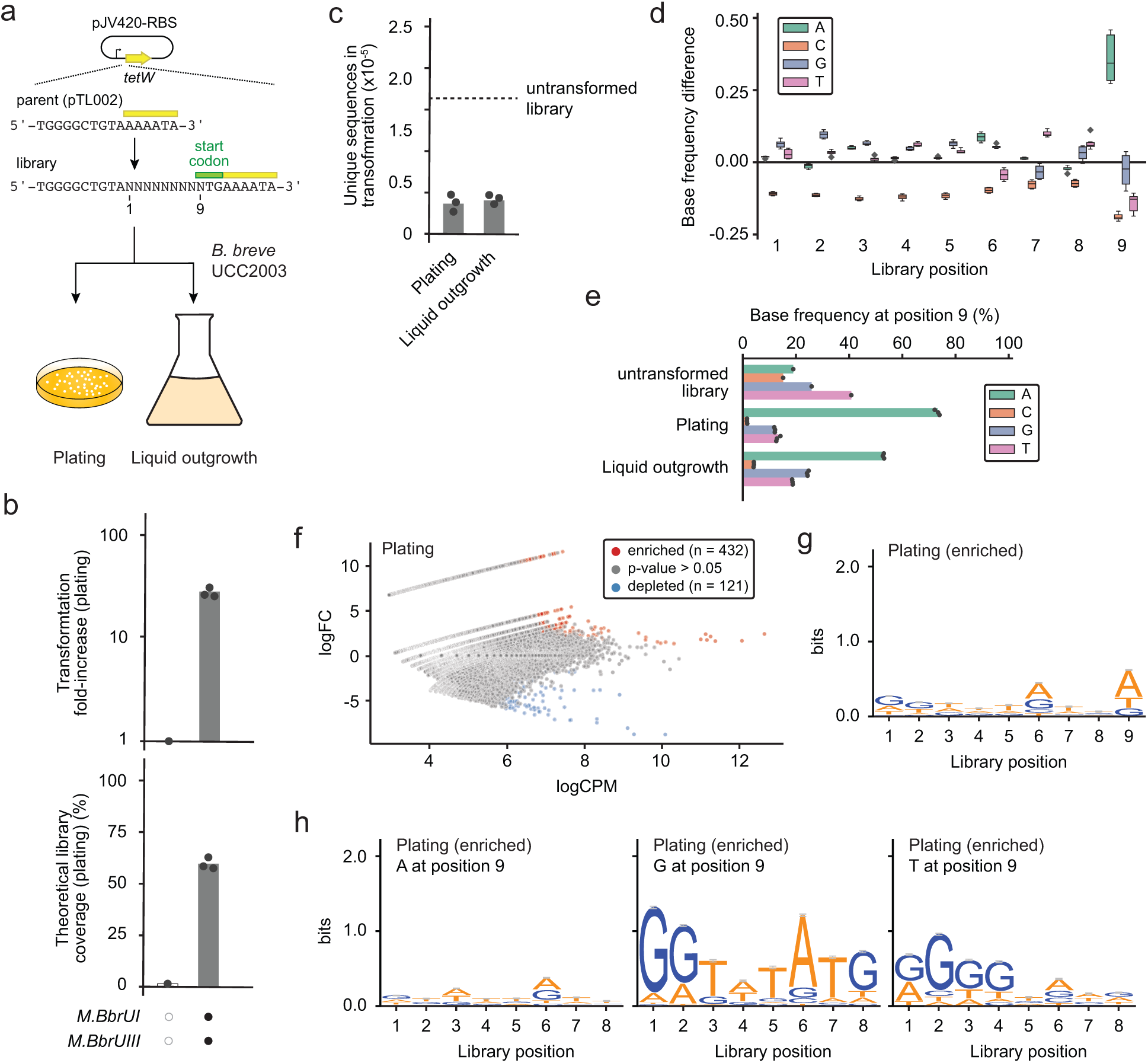
IMPRINT enhances a large library screen in *Bifidobacterium breve*. a, Overview of the library screen. The screen was conducted into two independent replicates with either direct plating followed by pooling the colonies or by liquid outgrowth prior to DNA extraction. **b,** Boost in library transformation with IMPRINT. **c,** Fraction of unique library sequences compared to the untransformed library following plating or liquid outgrowth after applying IMPRINT. See Table S4 for specific values. **d,** Difference in base frequencies at each library position compared to the untransformed library after applying IMPRINT. The box and whisker plots represent the combination of the three independent experiments from plating and the three independent experiments from liquid outgrowth after applying IMPRINT. **e,** Base frequencies at position 9 representing the first nucleotide of the original start codon. **f,** Enriched and depleted sequences within the transformed library after applying IMPRINT with plating. Each dot represents a specific sequence, and the plot captures the three independent replicates after applying IMPRINT with plating. See Methods for the determination of enriched or depleted sequences. **g,** Sequence logo for the enriched sequences within the transformed library after applying IMPRINT with plating. **h,** Sequence logo for the enriched sequences within the transformed library after applying IMPRINT with plating when base position 9 is an A (left), G (middle), or T (right). Letter sizes represent the enriched sequence conservation at that position after applying IMPRINT with plating. Error bars indicate an approximate Bayesian 95% confidence interval. Dots in b, c and e represent independent transformations followed by plating.

Assessment of the library sequenced via NGS revealed a number of unique features important for protein translation under our experimental setup (**Table S4**). First, we found that the number of unique sequences dropped from 62% of the library to an average of 14.6% (plating) or 16.2% (liquid outgrowth), indicating that a sizable fraction of the original library sequences could support sufficient *tetW* expression (**Fig. 6c**). The frequency of the parent vector also decreased an average of 93-fold in transformed samples, further indicating the loss of sequences that do not support translation (**Table S4**). Of the unique sequences in the plated transformants, the most represented nucleotide was an A as the first nucleotide of the start codon, as expected for the canonical start codon (**Fig. 6d-e**). G was also represented at this position, in line with GTG serving as an alternative start codon. Interestingly, T was represented more than C at this position, suggesting that Bifidobacteria can utilize TTG as an alternative start codon.

Within the set of sequences that supported transformation, we posited that sequences that were enriched within this set would reflect stronger expression of *tetW* and thus could provide unique insights into features governing translation. While identifying such sequences was complicated due to lower counts for each transformed sequence, we could identify 432 and 193 sequences that were significantly enriched (p < 0.05, n = 3) with plating (**Fig. 6f**) and liquid outgrowth (**Fig. S8a**), respectively. These sequences again contained an A, G, or T as the first nt in the start codon (i.e,, randomized position 9) (**Fig. 6g**), leading us to evaluate the enriched sequences associated with each start codon (**Figs. 6h and S8b-c**). For A at position 9, we did not observe any strong preferences, indicating that diverse sequences can drive sufficient *tetW* expression when paired with a canonical start codon. For G at position 9, an immediately upstream canonical start codon was enriched, suggesting that GTG was less sufficient in itself to drive translation. Finally, for T at position 9, there was no evidence of an immediately upstream canonical start codon; instead, G was strongly enriched 5-8 nts upstream of the TTG start codon, in line with a G-rich sequence helping drive translation. Therefore, coupling these three start codons with different RBS sequences could broadly tune the expression of heterologous genes in this recalcitrant bacterium. Overall, these results provide insights into rules governing translation in *B. breve* that became possible with the use of IMPRINT.

## DISCUSSION

Through this work, we demonstrated that IMPRINT offers a generalized means to recreate the pattern of DNA methylation in a host bacterium using TXTL, thereby boosting transformation of the methylated DNA into the host. As TXTL circumvents any need for cell culturing or protein purification required by traditional approaches, IMPRINT greatly accelerates the time from obtaining constructs for each MTase to achieving improved transformation. While plasmid constructs do require a cloning step that could prove challenging due to leaky expression of any cytotoxic MTases, we showed that MTases can be expressed from linear DNA templates that can be custom synthesized and used in IMPRINT shortly after receipt. Through IMPRINT, rapid methylation of DNA by simple and complex MTases, including type I MTases requiring multiple specificity genes, led to improved transformation efficiency in multiple bacterial species. Finally, applying HT-IMPRINT allowed the identification of the best MTase combination in a single transformation, simplifying subsequent transformation efforts.

While testing different MTases, we found that Type I MTases were less efficient at plasmid methylation, particularly in the presence of other MTases. We attributed the poor efficiency to the preference of many Type I MTases to methylate hemi-methylated DNA, which is not supplied as part of IMPRINT. While applying a Type I MTase separately from other MTases in sequential methylation reactions could compensate to some degree, we found that introducing mutations known to relax the preference for hemi-methylated DNA allowed the tested Type I MTase to efficiently methylate the supplied plasmid DNA. As the mutations can only be reasonably extended to related Type I MTases, future work could focus on identifying equivalent mutations in unrelated MTases across the broad diversity of Type I R-M systems found in nature^4,5,25,26^.

Outside of R-M systems, novel antiphage defense systems continue to be discovered throughout bacteria that could also impact DNA transformation^27,28^. While the vast majority of these systems appear to confer defense against bacteriophages, some have been shown to confer defense against conjugated plasmids^29^. Of these various systems, ones that rely on DNA methylation (e.g., BREX, DISARM)^30,31^ or other forms of DNA chemical modification could be incorporated into IMPRINT to allow transformed DNA to circumvent an even wider assortment of bacterial defenses. Regardless of the specific delivery mode or type of MTase, increased transformation by IMPRINT would open doors for efficient genetic manipulation and harnessing the rich diversity of bacteria for sustainable chemical production, enhancing food production, and applying cell-based biosensors and therapeutics^32,33^. IMPRINT could also be used to study the role of R-M systems in host defense and gene regulation, creating opportunities to explore the functional roles played by DNA MTases in diverse bacteria.

As a final demonstration of IMPRINT, we screened a large library of potential RBS and start-codon sequences driving expression of an antibiotic resistance gene in Bifidobacteria. The screen revealed a preference for a T as the first nucleotide of the start codon, suggesting that *B. breve* and possibly other Bifidobacteria can readily accommodate TTG as a start codon. TTG start codons are rare but utilized across the bacterial domain, with one comprehensive analysis concluding that 7.8% of genes initiate with this codon^34^. TTG is poorly utilized in *E. coli* though, with one study reporting a 97% reduction in gene expression when switching from ATG to TTG as the start codon^35^. The start codon heavily influenced which RBS sequences were enriched in *B. breve*, offering both the RBS and start codon as distinct means to tune gene expression in this probiotic bacterium.

### Limitations

While IMPRINT offers the opportunity to recreate methylation patterns and boost DNA transformation in diverse bacteria, there are notable limitations preventing the broad enhancement of transformation in any bacterium. First, IMPRINT currently relies on an *E. coli* cell lysate that functions optimally at 29℃; for hosts found in other environments such as high temperatures or salinities, their MTases would be unlikely to be functionally expressed in this TXTL reaction. However, TXTL systems derived from non-model bacteria and archaea are under development that could expand the conditions in which MTases can be expressed^36^. Another challenge is that IMPRINT is geared for transforming naked DNA. While this form of DNA is compatible with transformation procedures such as electroporation, chemical transformation, and natural transformation, as well as packaged into nanoparticles^37^, naked DNA is not readily compatible with cell-based delivery approaches such as conjugation or phage delivery. However, HT-IMPRINT could identify a minimal set of important MTases to express in the donor/packaging strain^38,39^. Finally, poor or even unsuccessful transformation could be attributed to a number of factors unrelated to R-M systems, such as poor transformation conditions (e.g., electroporator settings, growth state, recovery conditions), cell wall composition, and the ability of the transformed DNA to propagate and be selected. To assess whether R-M systems pose a major barrier, we recommend isolating and transforming plasmid DNA from the same strain if possible; a large boost in transformation would therefore indicate that IMPRINT could be successfully applied in this strain.

## METHODS

### Standard growth conditions

*E. coli* and *S. enterica* propagation was performed in LB medium (10 g/L NaCl, 5 g/L yeast extract, 10 g/L tryptone) while being shaken at 250 rpm at 37℃, aside from *E. coli* KL740 which was grown at 30℃. Plasmids were maintained at the following antibiotic concentrations: ampicillin (50 µg/mL), chloramphenicol (34 µg/mL).

Bifidobacteria were routinely grown in MRS liquid broth (BD CN# 288130) supplemented with 0.05% L-cysteine and MRS agar (BD CN# 288210) or Reinforced Clostridial Agar (RCA, Thermo CN# CM0151B) and incubated at 37℃ in an anaerobic chamber. Tetracycline was used to maintain plasmids at a concentration of 20 µg/mL.

### Generation of plasmid and linear DNA constructs

Plasmid pJV170 was used to readily clone new methyltransferases under P70a expression. Cloning was performed by amplifying plasmid pJV170 and host genomic DNA with primers that include homology tails, then by performing Gibson assembly according to manufacturer’s protocols (NEB CN# E2611S). The assembly mix was transformed into *E. coli* KL740 by electroporation, and colonies were screened using colony PCR and Sanger sequencing to determine if the clone was correct. Linear DNA constructs encoding Dam or eGFP were amplified via PCR from the plasmids pJV302 (Dam) and pJV170 (eGFP) using NEB’s Q5 HotStart High-Fidelity 2x Master Mix (NEB CN# M0494S). PCR reactions were treated with 1 μL of DPNI for 3 h at 37°C and purified with Zymo Research’s Clean & Concentrator-5 kit according to the manufacturer’s protocol (Zymo CN# D4004).

When necessary, small insertions or mutations were inserted into plasmids using Q5 site-directed mutagenesis (NEB CN# E0554S) according to manufacturer’s protocols and transforming chemically competent NEB 10-beta cells (NEB CN# C3019), where colony PCR and Sanger sequencing were again used to determine if the clone was correct.

To clone a library of barcoded *E. coli*-Bifidobacteria shuttle vectors, Q5 mutagenesis was performed according to manufacturer’s protocols where four random nucleotides were added to the 5’ end of one of the primers. The Q5 mutagenesis mix was transformed into NEB 10-beta chemically competent cells, and individual barcodes were isolated using colony PCR and Sanger sequencing to determine the barcode sequence of each clone.

The current RBS and the start codon were deleted from pJV420 by performing Q5 mutagenesis to generate the parent plasmid pTL002 for RBS library construction. The backbone of the library was amplified from pTL002 with homology tails by Q5 Hot Start High-Fidelity 2X Master Mix (NEB CN# M0494S). 125-nt ssDNA with 9 random nucleotides at the RBS and the first base of the start codon region was synthesized by Eurofins. After annealing at 65℃ for 30s with a primer on the 125-nt ssDNA end, a single-cycle extension with NEB Ultra Q5 II master mix was performed at 72℃ for 2 min to make a double-stranded insert. RBS library backbone and insert were assembled by Gibson assembly and transformed into *E. coli* NEB10β (C3020K) by electroporation. 20 µL of recovered cell culture was plated to obtain colonies for calculating transformation efficiency and performing colony PCR and Sanger sequencing to initially confirm cloning of the RBS library. The rest of the recovered culture was cultured in LB with ampicillin at 25℃ and 250 rpm until reaching an OD_600_ of ∼0.6. RBS library plasmids were extracted by ZymoPURE II Plasmid Midiprep Kit (D4200) and further propagated in methyltransferase-deficient *E. coli* EC135 to yield unmethylated RBS library plasmids.

### *In vitro* methylation of plasmid and linear DNA

To methylate plasmid and linear DNA *in vitro* using IMPRINT, TXTL reactions were first set up by incubating myTXTL Sigma 70 master mix (Arbor Biosciences CN# 507024) or myTXTL Linear DNA Expression Kit (Arbor Biosciences CN#508024) with each methyltransferase plasmid or linear DNA harboring the methyltransferase gene(s) at 10 nM to be expressed at 29℃ for 12-16 hours. The shuttle plasmid was first propagated in methyltransferase-deficient *E. coli* EC135 to yield an unmethylated shuttle plasmid. Unmethylated linear DNA was generated by PCR using NEB’s Q5 HotStart High-Fidelity 2x Master Mix (NEB CN# M0494S) from plasmid extracted from *E. coli* DH5a, treated with 1 μL of DpnI for 3 h at 37℃ and purified with Zymo Research’s Clean & Concentrator-5 kit according to the manufacturer’s instructions (Zymo CN# D4004). Then, a methylation reaction was set up (50 µL) by mixing a total of 1 µL of the TXTL reaction(s) with 1 - 2 μg of the shuttle vector or 1 μg of unmethylated linear DNA, 1x Dam methyltransferase buffer (NEB CN# M0222) and 640 µM AdoMet (NEB CN# B9003). The methylation reactions were incubated at 37℃ for 1-4 h. Following methylation, the reaction was first treated with 100 µg/mL Proteinase K and incubated at 50℃ for 30 min then with 100 µg/mL RNase A and incubated at 37℃ for 1 h to remove impurities from the TXTL mix. Finally, the shuttle plasmid was purified using column purification (Zymo Research CN# D4013) according to manufacturer protocols.

### Electroporation

Electroporation of *S. enterica* was performed as follows: an overnight culture was diluted 1:20 in fresh LB media and grown until an OD_600_ of ∼0.6. Then, cells were harvested by centrifugation at 5,000 rpm and 4℃, before being washed twice with 25 mL of ice-cold 10% glycerol. Harvested cells were then suspended in 0.5 mL ice-cold 10% glycerol. 0.1 mL of cell suspension was added to 1-mm electroporation cuvettes with 50-100 ng of plasmid DNA, and the cells were electroporated at 1.8 kV, 200 Ω resistance, and 25 µF. 0.9 mL of pre-warmed LB broth was then added to the electroporated cells and a recovery was set up for 1 h at 37℃ before cells were plated on LB agar supplemented with chloramphenicol. Colonies were counted after overnight incubation at 37℃.

To transform *B. breve* and *B. longum* by electroporation, cells were first made electrocompetent by adapting a previously published protocol^40^. In short, 4 mL of an overnight culture was added to 50 mL of mMRS supplemented with 1% glucose and 0.05% L-cysteine, and cells were grown at 37℃ in an anaerobic chamber until the OD_600_ reached 0.6 (*B. breve*) or 1.0 (*B. longum*). Then, cells were harvested by centrifugation at 4,500 rpm and 4℃, and washed twice with 25 mL of ice-cold wash buffer (0.5 M sucrose, 1 mM ammonium citrate, pH 6.0) then once more with 1 mL of ice-cold wash buffer before resuspending the cells in 0.375 mL ice-cold wash buffer. To perform electroporation, 90 µL of the cell suspension was added to 1-mm electroporation cuvettes with 200-500 ng of plasmid DNA, and electroporation was performed at 2.0 kV, 200 Ω resistance, and 25 µF. For *B. longum*, the cells were incubated with the plasmid DNA on ice for at least 5 min before performing electroporation. After electroporation, 0.9 mL of pre-warmed Reinforced Clostridial Medium (RCM) was added to the cells and a recovery was set up for 3 h at 37℃ in an anaerobic chamber. Finally, cells were plated on Reinforced Clostridial Agar (RCA) supplemented with tetracycline, and colonies were counted after 2-3 days of incubation at 37℃.

### HT-IMPRINT

To determine the optimal methylation pattern required for transformation of different Bifidobacterial strains, 4-nt barcodes were cloned into the *E. coli*-*Bifidobacterium* shuttle plasmid (see Tables S2 and S3 for barcodes). Then, 2-3 barcodes were assigned to each combination of DNA MTases for a given strain. IMPRINT reactions were performed to methylate the barcoded shuttle plasmids with each combination of the methyltransferases present in the strain. The purified, barcoded shuttle vectors from the IMPRINT reactions were then pooled together and transformed into the Bifidobacterial strain, and cells were both plated on RCA supplemented with tetracycline and diluted in MRS broth supplemented with 0.05% L-cysteine and tetracycline. After 2-3 days, the transformed plasmids were purified from the back-diluted cultures, or from the grown colonies by first adding 1 mL PBS to the agar plates to resuspend colonies. In both cases, the plasmid was used as a template to amplify a segment of the plasmid harboring the barcode. The PCR amplicon was submitted for Amplicon-EZ Sequencing performed by Genewiz (https://www.genewiz.com/en/Public/Services/Next-Generation-Sequencing/Amplicon-Sequencing-Services/Amplicon-EZ). The number of reads mapping to each barcode was counted within each .fastq.gz file using the command zgrep -c “CTGCNNNN” *.fastq.gz for pJV420_U barcodes or zgrep -c “GCTTNNNN” *.fastq.gz for pJV420_D barcodes. From there, the median barcode count (if using three barcodes per sample) or average barcode count (if using two barcodes per sample) was calculated for each methylation sample in the transformed cells and in the pool control. The ratio of each sample in the transformed cells compared to the pool control was compared to the ratio of total reads in the transformed cells relative to the pool control to calculate the extent of barcode enrichment or depletion.

### Analysis of the RBS library

Paired-end sequencing FASTQ files were merged using BBmerge (version 38.69) with parameters “qtrim2=t ecco trimq=20 -Xmx1g mix”. An in-house Python script was used to identify and count the 9-nt sequences, including the 8-nt putative RBS sequence and first nt of the start codon, based on the flanking sequences in the plasmid. The enrichment analysis was performed separately for plating and liquid culture samples. After filtering for at least 10 counts per million in 2 samples, log-fold change (logFC) of the sequences between the transformants and original library was estimated using edgeR (version 3.28.1) and a quasi-likelihood F-test was used to test for significance after fitting in a generalized linear model. Sequences with a logFC higher than 0 and a P-value lower than 0.05 were defined as enriched. Enriched patterns in sequences were visualized using WebLogo (version 3.7.12). Input FASTA files for WebLogo included N times of the sequences (N = the mean counts in three replicates for enriched sequences), while the nucleotide composition was calculated based on the original library.

## Data Availability

The NGS data from HT-IMPRINT is available through NCBI GEO GSE189864, with an access token for peer review of ufiligswxpwtbwb. The NGS data from the RBS library experiment is available through NCBI GEO GSE240651, with an access token for peer review of qfkdqwcornyltyv.

## Statistical analyses

A student’s t-test was utilized to determine statistical significance in DNA transformation from different MTases. In all cases, the t-test was a two-sample, equal variance dataset with a two-tailed distribution. Two-sided Mann-Whitney U test was used to determine the statistical significance in the difference of AG count between enriched and depleted RBS sequences.

## Acknowledgements

We thank Douwe Sinderen for providing the Bifidobacteria-*E. coli* shuttle vector pJV420 and the strain *Bifidobacterium breve* UCC2003. Frank Englert, Tatjana Achmedov, and Isabell Köblitz assisted with next-generation sequencing of the RBS library. We also thank Tingyi Wen for providing *E. coli* strain EC135. This work was supported through funding by the National Science Foundation (MCB-1452902 to C.L.B.; CBET-1934284, EFMA-2029327, IOS-2120593 to N.C.), the Deutsche Forschungsgemeinschaft (BE 6703/2-1 to C.L.B.), the Novo Nordisk Foundation (NNF19SA0035474 to N.C.), the Camille & Henry Dreyfus Foundation (2017-137 to C.L.B.), and the Bavarian State Ministry for Science and the Arts through the research network bayresq.net (to L.B.).

## Author Information

J.M.V. and C.L.B. conceptualized the IMPRINT method, designed the experiments, analyzed the data, and wrote the manuscript. J.M.V., S.S., F.T., and J.V.S. developed and applied IMPRINT in *E. coli* and *S. enterica*. J.M.V., D.D., and T.L. applied IMPRINT and developed HT-IMPRINT in Bifidobacteria. N.C. provided critical feedback on experimental design and contributed to scientific discussions around developing HT-IMPRINT. T.L. generated the RBS library and performed the library screen. Y.Y. and L.B. analyzed the RBS library. Y.Y. D.D., S.S., J.V.S., Y.Y., L.B., and N.C. commented on the manuscript.

## Ethics Declarations

J.M.V. and C.L.B. have filed a provisional patent application related to this work. C.L.B. is a co-founder and Scientific Advisory Board member of Locus Biosciences as well as a Scientific Advisory Board member of Benson Hill. The other authors declare no competing interests.

