## Supplementary Information for "A cell-free transcription-translation pipeline for recreating methylation patterns boosts DNA transformation in bacteria"

### SUPPLEMENTARY FIGURES

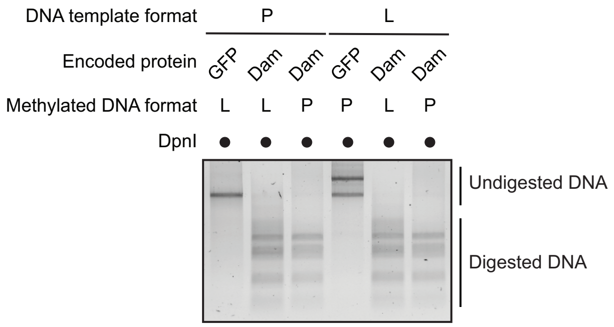

**Supplementary Fig. S1: M.EcoKdam expressed from a plasmid or linear DNA template can drive methylation of plasmid and linear DNA**. Four separate TXTL reactions using myTXTL Linear DNA Expression Kit were performed with two plasmid templates encoding either M.EcoKdam (pJV302) and GFP (pJV170) and two PCR-derived linear DNA templates from these plasmids containing the P70a promoter, a gene encoding M.EcoKdam or GFP, and a rho-independent terminator. Methylation reactions were then performed from the overnight TXTL reactions by adding plasmid pMRTK-20sfGFP or linearized pMRTK-20sfGFP, a commercial Dam methylation buffer, and methyl donor SAM. The plasmid and linear DNA were then purified from the reaction, and DpnI restriction digestion was performed to assess whether the plasmid was Dam-methylated.

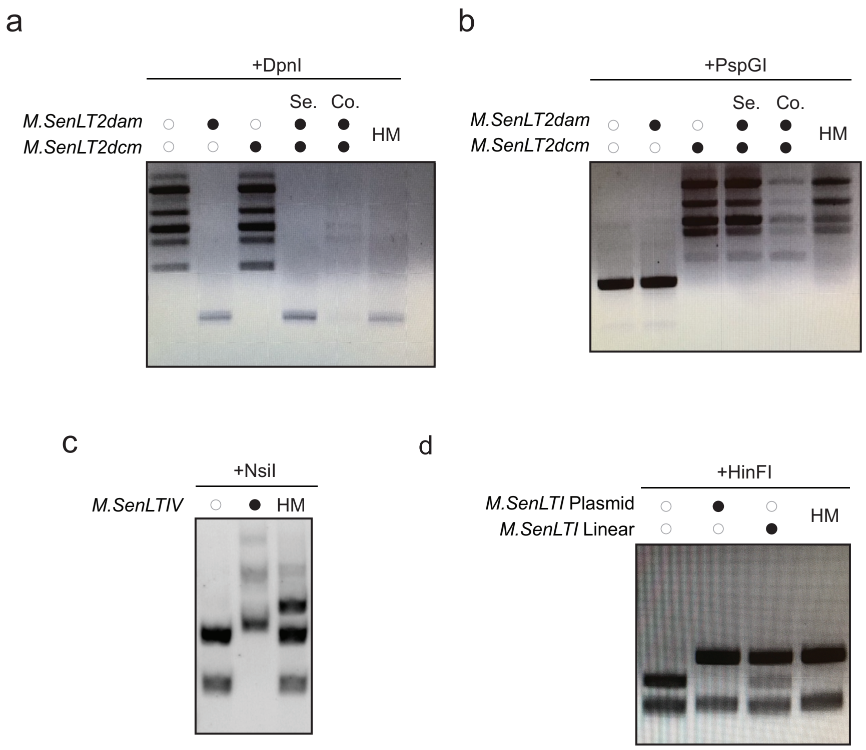

**Supplementary Fig. S2: Validation of methylation by *S. enterica* LT2 methyltransferases. a-b,** M.SenLT2dam and M.SenLT2dcm methylation. To validate that IMPRINT could be used to methylate plasmids with methyltransferases from *S. enterica* LT2, M.SenLT2dam and M.SenLT2dcm were both expressed on plasmids, either in the same TXTL reaction (Co.) or in separate TXTL reactions (Se.). Methylation reactions were then set up with either one or both methyltransferases using plasmid pJV400. After plasmid cleanup, DpnI (**a**) or PspGI (**b**) restriction digestions were performed to test if the plasmid was either Dam- or Dcm-methylated, respectively. **c,** Methylation by orphan methyltransferase M.SenLT2IV. An IMPRINT expression construct harboring M.SenLT2IV was added to a TXTL reaction. Then, a methylation reaction was performed with plasmid pJV447 that harbored two NsiI/M.SenLT2IV motifs. Methylation was assessed by restriction digestion with NsiI that would not cleave DNA methylated by M.SenLT2IV. **d,** Methylation by Type III methyltransferase M.SenLT2I. M.SenLT2I was expressed in TXTL either by cloning the methyltransferase into an IMPRINT expression construct or by amplifying it from *S. enterica* LT2 genomic DNA using primers that included T7 promoter and T500 terminator sequences. A T7 RNA polymerase expression plasmid pJV441 was added to the linear expression reaction along with the GamS protein to protect the DNA from degradation by RecBCD present in the TXTL lysate. The methyltransferase reactions were used to methylate the plasmid pJV184 harboring an overlapping HinFI/M.SenLT2I motif. After plasmid purification, restriction digestion with HinFI was performed, which would cut plasmid pJV184 seven times if not methylated by M.SenLT2I but only six times if methylated by M.SenLT2I. HM: *S. enterica* LT2 host-methylated plasmid.

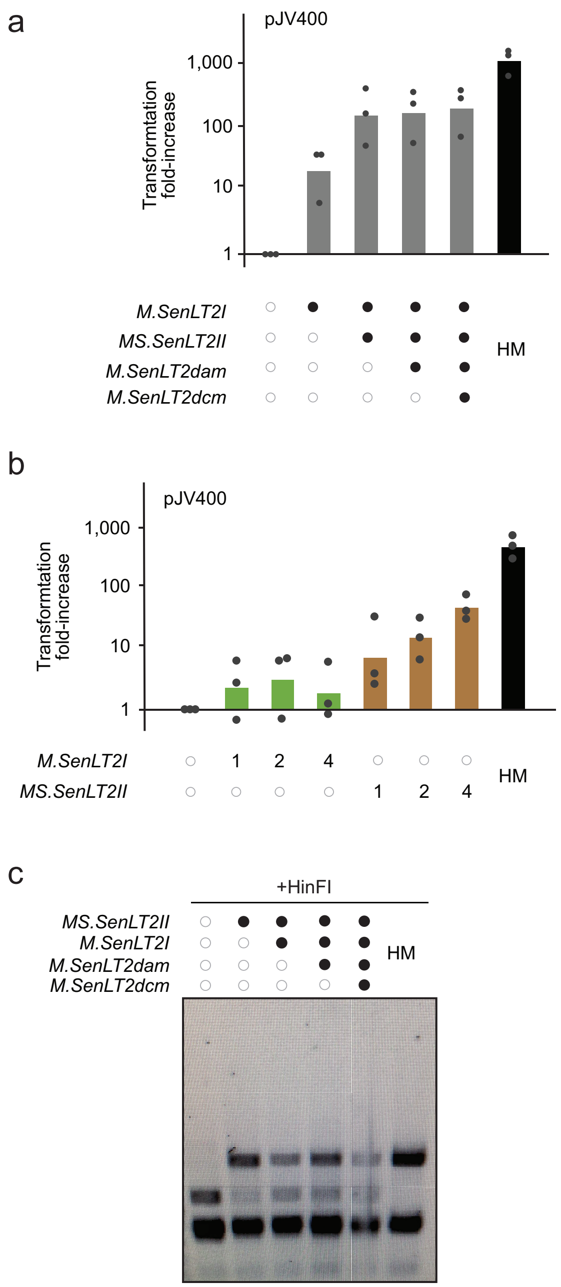

**Supplementary Fig. S3: Incomplete methylation by M.SenLT2II. a,** Series methylation of methyltransferases in *S. enterica* LT2. To overcome poor multiplexed methylation, a series methylation reaction was set up where shuttle plasmid pJV400 was first methylated with MS.SenLT2II, then the plasmid was purified and a second IMPRINT reaction was performed to methylate pJV400 with remaining methyltransferases M.SenLT2I, M.SenLT2dam*,* and/or M.SenLT2dcm. The methylated plasmids were then transformed into *S. enterica* LT2, where fold change in CFU relative to unmethylated pJV400 was determined as well as a protection score relative to unmethylated and host-methylated (HM) pJV400 controls. Dots represent transformations from three separate IMPRINT reactions, and the bar represents the average fold change in CFU as well as the protection score. **b,** Time course experiment with M.SenLT2I and MS.SenLT2II. To assess how quickly complete methylation was achieved by R-M methyltransferases in *S. enterica* LT2, methylation reactions of 1-, 2-, or 4-hours were performed using either M.SenLT2I or MS.SenLT2II and shuttle plasmid pJV400. Methylated plasmids were then transformed into *S. enterica* LT2 along with unmethylated pJV400 and host-methylated pJV400 (HM). A fold change relative to unmethylated pJV400 was calculated, and a protection score was determined for each methylated plasmid relative to unmethylated and host-methylated controls. Dots represent transformations from three separate IMPRINT reactions, and the bar represents the average fold change in CFU as well as the protection score. **c,** Methylation by MS.SenLT2II is incomplete. In order to determine if MS.SenLT2II was completely methylating plasmid DNA, plasmid pJV412 harboring an overlapping HinFI/MS.SenLT2II motif was methylated with MS.SenLT2II alone or in combination with other *S. enterica* LT2 methyltransferases M.SenLT2I, M.SenLT2dam, and M.SenLT2dcm. After the plasmid was purified from the IMPRINT reactions, a HinFI restriction digestion was performed, which would cleave pJV412 seven times if the plasmid was not methylated by MS.SenLT2II and would only cleave pJV412 six times if the plasmid was methylated by MS.SenLT2II.

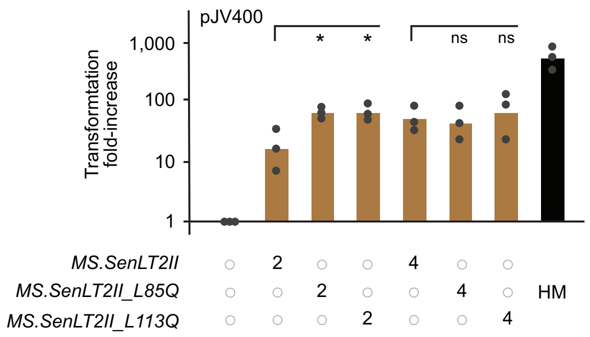

**Supplementary Fig. S4: Time-course experiment comparing methylation by MS.SenLT2II and the mutants MS.SenLT2II_L85Q and MS.SenLT2II_L113R**. To assess how quickly complete methylation was achieved by type I R-M methyltransferase MS.SenLT2II and MS.SenLT2II_L85Q and MS.SenLT2II_L113R, methylation reactions of 2 hours or 4 hours were performed by combining each methyltransferase with shuttle plasmid pJV400. Methylated plasmids were then transformed into *S. enterica* LT2 along with unmethylated pJV400 and host-methylated pJV400 (HM). A fold change relative to unmethylated pJV400 was calculated, and a protection score was determined for each methylated plasmid relative to unmethylated and host-methylated controls. Dots represent transformations from three separate IMPRINT reactions, and the bar represents the average fold change in CFU as well as the protection score. The bottom of the bar marks the reference for statistical analysis. **: $p < 0.01$. *: $p < 0.05$. ns: $p > 0.05$.

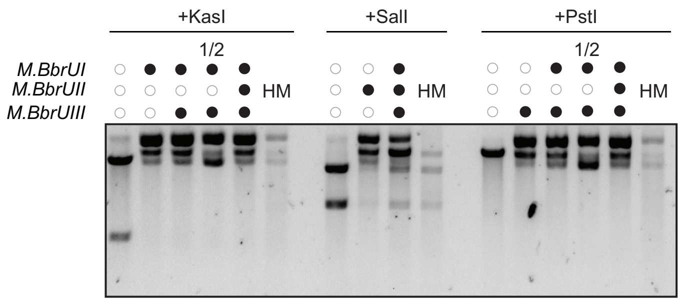

**Supplementary Fig. S5: IMPRINT methylates a shuttle plasmid with all three R-M methyltransferases from *B. breve* UCC2003**. To assess whether IMPRINT could be used to methylate plasmids with methyltransferases from Bifidobacteria, three type III R-M methyltransferases from *B. breve* UCC2003 (M.BbrUI, M.BbrUII, and M.BbrUIII) were expressed in separate TXTL reactions and added either separately or in combination to methylate shuttle plasmid pJV420. For one sample, half the amount of methyltransferases were added to the methylation reaction to determine if this would improve plasmid cleanup from TXTL lysate without affecting methylation ("½" sample). Methylation by M.BbrUI was validated by restriction digestion of plasmid pJV420 with KasI which would only cleave DNA not methylated by M.BbrUI. Methylation by M.BbrUII was validated by restriction digestion of plasmid pJV420 with SalI which would only cleave DNA not methylated by M.BbrUII. Methylation by M.BbrUIII was validated by restriction digestion of plasmid pJV420 with PstI which would only cleave DNA not methylated by M.BbrUIII. HM = host-methylated (pJV420 propagated in *B. breve* UCC2003).

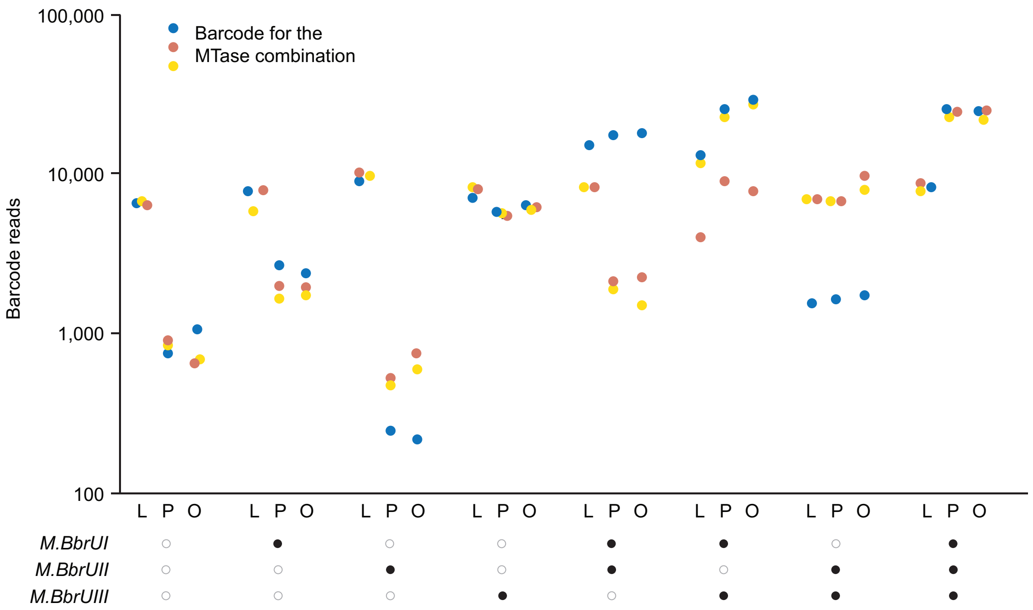

**Supplementary Fig. S6: Barcode abundances when performing HT-IMPRINT in *B. breve* UCC2003.** The 8 possible methylation patterns from *B. breve* UCC2003 were assigned three barcodes each on pJV420_D (blue = barcode 1, yellow = barcode 2, red = barcode 3), and the pool of methylated plasmids were transformed together into *B. breve* UCC2003. Amplicons containing the barcoded portion of pJV420_D from the pooled library prior to transformation (L), the library from the plated colonies (P), and the library from the liquid outgrowth (O) were sent for high-throughput amplicon sequencing, where the number of reads from each barcode was determined.

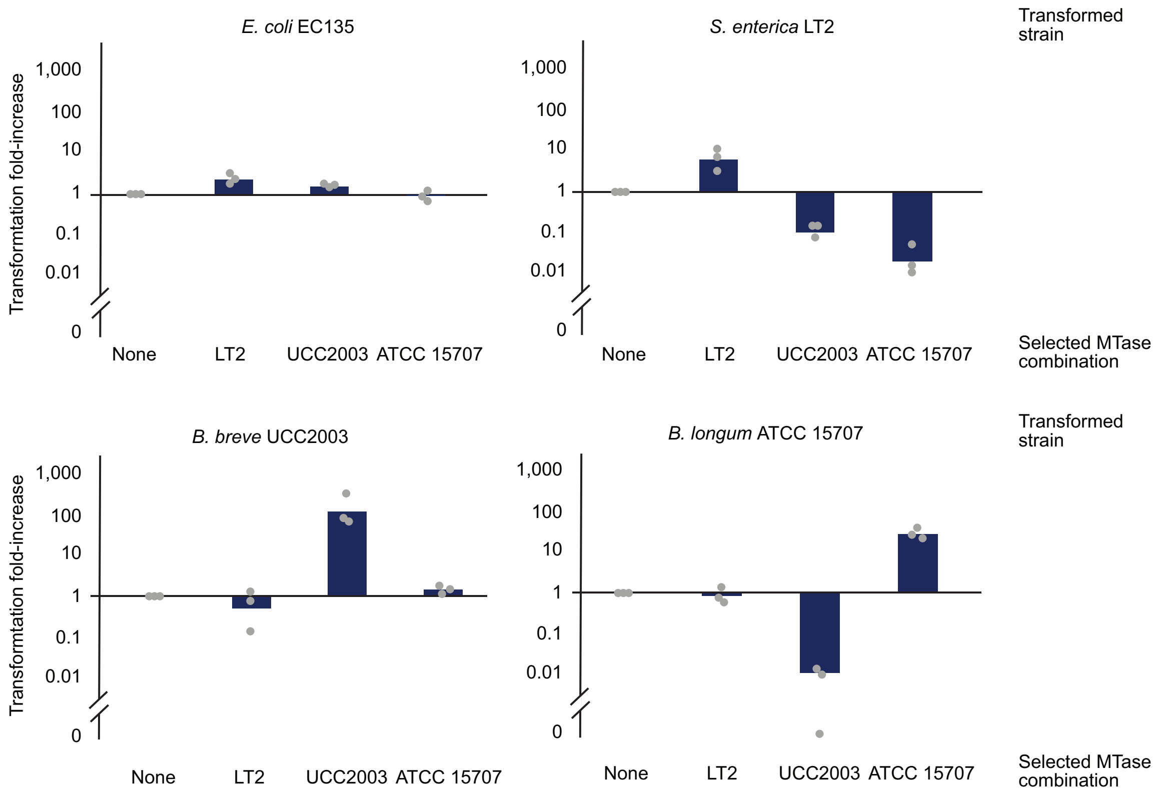

**Supplementary Fig. S7: Cross-transformations of optimally methylated plasmids from different strains**. To determine whether the optimal methylation pattern determined from each tested strain (*S. enterica* LT2, *B. breve* UCC2003, and *B. longum* ATCC15707) was specific to that strain only, a cross-transformation experiment was performed where shuttle plasmid pJV420 harboring each tested strain's optimal methylation pattern was used to transform the other strains. Plasmids that were unmethylated, methylated by MS.SenLT2II_L85Q and M.SenLT2I from *S. enterica* LT2, methylated by M.BbrUI and M.BbrUIII from *B. breve* UCC2003, or methylated by $M_2$.Blo1217ORF1038 and $MS_2$.Blo1217ORF1481 from *B. longum* ATCC15707 were transformed into each strain, where *E. coli* EC135 was used as a control strain lacking R-M systems. The fold change in CFU was determined for each methylated sample relative to unmethylated pJV420. Dots represent the fold changes obtained from three individual transformations, and bars represent the average fold change.

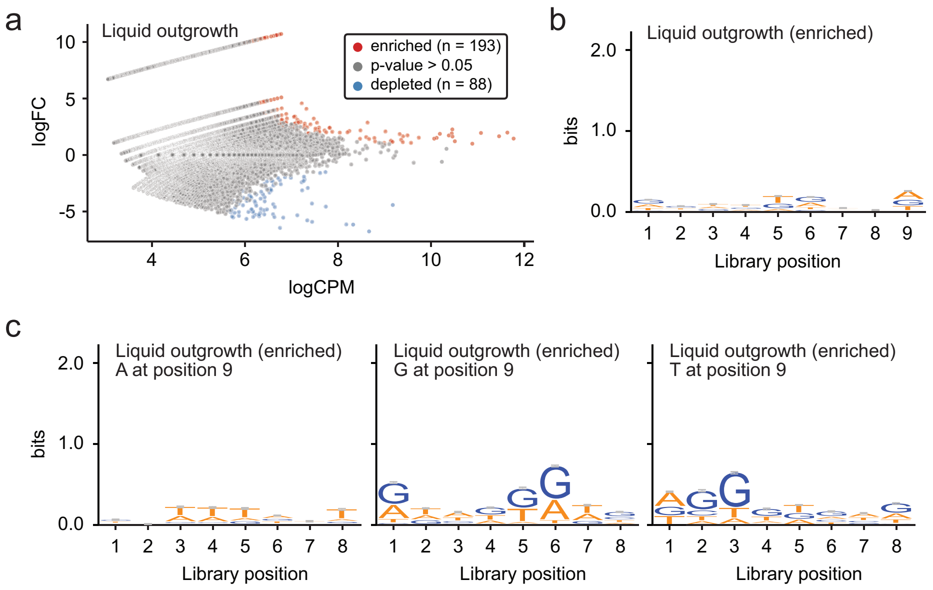

**Supplementary Fig. S8: Assessment of enriched RBS sequences with liquid outgrowth**. **a,** Enriched and depleted sequences within the transformed library after applying IMPRINT with liquid outgrowth. Each dot represents a specific sequence, and the plot captures the three independent replicates after applying IMPRINT with liquid outgrowth. See Methods for the determination of enriched or depleted sequences. **b,** Sequence logo for the enriched sequences within the transformed library after applying IMPRINT with liquid outgrowth. **c,** Sequence logo for the enriched sequences within the transformed library after applying IMPRINT with liquid outgrowth when base position 9 is an A (left), G (middle), or T (right). Letter sizes represent the enriched sequence conservation at that position after applying IMPRINT with liquid outgrowth. Error bars indicate an approximate Bayesian 95% confidence interval.

### SUPPLEMENTARY TABLES

**Supplementary Table S1**: Comparison of IMPRINT with other methods of bypassing R-M barriers to DNA transformation. Different methods of overcoming host R-M systems to boost transformation are compared, including plasmid artificial modification (PAM)[1], methods involving methylation *in vitro* with purified methyltransferases or in a host cell lysate[2,3], SyngenicDNA[4], and methods that involve deleting restriction endonuclease genes associated with R-M systems[5]. Methods were compared based on how many R-M systems they could reasonably overcome, DNA compatibility, the average time the method takes to set up for a new strain, and any associated toxicity issues or other issues impacting their broad use.

| **Method** | IMPRINT | PAM | Purified MTases | Cell lysate | SyngenicDNA | KO R-M |
|---|---|---|---|---|---|---|
| **Max. # of MTases** | >10 | 1-2 | 1-2 | >10 | >10 | 1-2 |
| **DNA Compatibility restrictions** | None | Must replicate in *E. coli* and strain-of-interest, only circular DNA | None | Only circular DNA | Must remove R-M motifs from DNA | None |
| **Speed*** | < 1 day | Days to weeks (starting from MTase expression constructs) | Weeks with purification (starting from MTase expression constructs) | < 1 day | Months (Methylome sequencing followed by DNA synthesis, cloning) | Weeks to months (genome editing) |
| **Toxicity issues** | None | MTase toxicity in *E. coli* | MTase toxicity during overexpressi on | None | None | Deletion can impact strain physiology |
| **Other issues** | None | None | Limited to-date to Type II MTases | Inefficient methylation | Requires bioinformatics pipeline | Need for available genetic tools |

*Timing begins when DNA constructs for expressing MTases are available. No MTase constructs are needed for SyngenicDNA and KO R-M, so these start with the initial experimental procedures (methylome sequence and generating gene knockouts, respectively).

**Supplementary Table S2:** List of barcodes used for HT-IMPRINT in *B. breve* UCC2003.

| Barcode Numbers | Barcode Sequences | Methylation Combination |
|---|---|---|
| D1-D2-D3 | GTGG-GAAG-GCTG | Empty |
| D4-D5-D6 | GGAG-ACGA-TCGA | *M.BbrUI* |
| D7-D8-D9 | CACA-GCAC-TAAA | *M.BbrUII* |
| D10-D11-D12 | CCCG-CCAT-ACAT | *M.BbrUIII* |
| D13-D14-D15 | CGTC-CAAG-GGAA | *M.BbrUI* + *M.BbrUII* |
| D16-D17-D18 | AAAG-TGCC-GGAT | *M.BbrUI* + *M.BbrUIII* |
| D19-D20-D21 | AAAC-TGAA-CTCG | *M.BbrUII* + *M. BbrUIII* |
| D22-D23-D24 | TGTC-CTGA-CGCC | *M. BbrUI* + *M.BbrUII* + *M. BbrUIII* |

**Supplementary Table S3:** List of barcodes used for HT-IMPRINT in *B. longum* ATCC15707.

| Barcode Numbers | Barcode Sequences | Methylation Combination |
|---|---|---|
| U1-D1 | ATCT-GTGG | Empty |
| U2-D2 | CGTT-GAAG | *MS2.Blo1217ORF1481* |
| U3-D3 | TCGT-GCTG | *M1.Blo1217ORF1038* |
| U4-D4 | ACTT-GGAG | *M2.Blo1217ORF1038* |
| U5-D5 | AATA-ACGA | *MS1S2.Blo1217ORF1481* |
| U6-D6 | GTGT-TCGA | *MS2.Blo1217ORF1481 + M1.Blo1217ORF1038* |
| U7-D7 | TCTA-CACA | *MS2.Blo1217ORF1481+ M2.Blo1217ORF1038* |
| U8-D8 | CCTC-GCAC | *M1M2.Blo1217ORF1038* |
| U9-D9 | GGTT-TAAA | *MS1S2.Blo1217ORF1481 + S1.Blo1217ORF1481* |
| U10-D10 | CGCA-CCCG | *MS1S2.Blo1217ORF1481+ M2.Blo1217ORF1038* |
| U11-D11 | CTTA-CCAT | *MS2.Blo1217ORF1481 + M1M2.Blo1217ORF1038* |
| U12-D12 | TTAG-ACAT | *MS1S2.Blo1217ORF1481 + M1M2.Blo1217ORF1038* |

**Supplementary Table S4**: Read counts from RBS library screen in *B. breve* UCC2003.

| Sample | Total reads | Unmapped reads (%) | Parent reads | Mapped ibrary reads | Mapped ibrary reads (%) | Unique library reads |
|---|---|---|---|---|---|---|
| Untransformed library | 2852009 | 2.1514 | 1553152 | 1185403 | 42.49 | 163721 |
| Plating_biorep 1 | 2307277 | 3.3946 | 14657 | 2172960 | 97.49 | 27415 |
| Plating_biorep 2 | 2961096 | 2.7413 | 13277 | 2823181 | 98.03 | 48375 |
| Plating_biorep 3 | 2649802 | 2.8826 | 11073 | 2521179 | 97.97 | 38782 |
| Outgrowth_biorep1 | 2114495 | 2.8454 | 23161 | 1986890 | 96.72 | 47411 |
| Outgrowth_biorep2 | 2380682 | 4.0741 | 24406 | 2184212 | 95.64 | 43945 |
| Outgrowth_biorep3 | 3196711 | 2.8607 | 24286 | 3021924 | 97.32 | 36542 |

**Supplementary Table S5:** Plasmids used in this study.

| Plasmid | Description | Plasmid map |
|---|---|---|
| pJV170 | P70a-GFP plasmid to clone methyltransferases | https://benchling.com/s/seq-QKS9eCV32deK7RTaY4iX?m=slm-whD895YIrMKPbOOScN1I |
| pJV441 | P70a-T7 RNA polymerase expression construct | https://benchling.com/s/seq-BVJEFTHnKix0NxHtCqF5?m=slm-JfM0yu4RGmDL521e3oVq |
| pJV24 | *M.EcoKdam* and *M.EcoKdcm* validation | https://benchling.com/s/seq-LBM9s4EYOtkBiERMJrnD?m=slm-bG8trK8GcJHMgWm1ZLN6 |
| pMRTK-20sfGFP | *M.EcoKdam* linear expression validation | https://benchling.com/s/seq-0dFxPHU3XtxdLge0jyAX?m=slm-LCIITt1RPcRmBtsIINVR |
| pJV400 | LT2 shuttle vector ColE1 origin | https://benchling.com/s/seq-jfs3oAJOBfy2Bc8HP4cR?m=slm-jtyKyagVktbSNtsY3e0c |
| pJV414 | LT2 shuttle vector p15A origin | https://benchling.com/s/seq-p4ST6tBYAVdcSKkQ46wE?m=slm-a5ysOUDUujoscjrEyAug |
| pJV420 | Bifidobacteria-*E. coli* shuttle vector | https://benchling.com/s/seq-mvth5uDmtTwQVcRWen6X?m=slm-Hrq3dwA61YnoAeuDXZji |
| pJV302 | *M.EcoKdam* expression construct | https://benchling.com/s/seq-zIRYJPvWpX0m1VL88ybI?m=slm-MsaVRc9GH8xDsrbu44sh |
| pJV593 | *M.EcoKdcm* expression construct | https://benchling.com/s/seq-CZIWHrWAIqPCZOiO5fOO?m=slm-HWkasz34xucCDAVjcn1J |
| pJV388 | *M.SenLT2dam* expression construct | https://benchling.com/s/seq-YHeAIUMK2jmo3kdQw1Gs?m=slm-uLrEOWFutqrJfp8FVQfP |
| pJV389 | *M.SenLT2dcm* expression construct | https://benchling.com/s/seq-MhdR9t1yxUzsLfyQZLTR?m=slm-OWUZft3LhTMAt1wLOXR5 |
| pJV303 | *M.SenLT2I* expression construct | https://benchling.com/s/seq- |

| | | |
|---|---|---|
| | | 7zdYwfROPjYXRXIcGDZa?m=slm-ggslC6gkpkxxhjNNymHb |
| pJV265 | *MS.SenLT2II* expression construct | https://benchling.com/s/seq-s5trithkW8Z9hQ36YKiR?m=slm-Es2oD8onPFUfNBAw0faT |
| pJV505 | *MS.SenLT2II_L85Q* expression construct | https://benchling.com/s/seq-SJQsWVAfyHmTNpSrXfz7?m=slm-J628G3xsYNqVOBcODYd6 |
| pJV506 | *MS.SenLT2II_L113R* expression construct | https://benchling.com/s/seq-lQRAmjcvHlTuGsnPcVPU?m=slm-uoGuOdLibfLdnUk5VjxE |
| pJV354 | *M.SenLT2IV* expression construct | https://benchling.com/s/seq-3AUPzdlUI6ZdfvE7k3ZA?m=slm-N1PYpz5Hu8CSAM9sZUiO |
| pJV184 | *M.SenLT2I* validation plasmid | https://benchling.com/s/seq-3hL0DbE5d6y0aj6j01Sn?m=slm-pEslkMAGvUlpVLloKoDg |
| pJV412 | *MS.SenLT2II* validation plasmid | https://benchling.com/s/seq-t3ioJvLnqn78yGxFglks?m=slm-XMAzWaAo6CRRyG1XCPS2 |
| pJV447 | *M.SenLT2IV* validation plasmid | https://benchling.com/s/seq-egWA09ByPTMeBa60xTPq?m=slm-G0kVaR8frDgtXnLAlspd |
| pJV457 | *M.BbrUI* expression construct | https://benchling.com/s/seq-oTIPpDXWTbC3t6eymOah?m=slm-rv8Di1WfiCZGwe8R1tzc |
| pJV476 | *M.BbrUII* expression construct | https://benchling.com/s/seq-keb6VaRGtou3cU0Fthtr?m=slm-y9XVvvDfPIYhtgRWUv6O |
| pJV461 | *M.BbrUIII* expression construct | https://benchling.com/s/seq-XhzqE9uH4FNkngCv4RC3?m=slm-Blpa3SGxKPe4mY39jNle |
| pJV544 | *M1.Blo1217ORF1038* expression construct | https://benchling.com/s/seq-l0CE0ukWudaJ9pLOefem?m=slm-Kwc8mWF9jOAWUUxeGAYM |
| pJV466 | *M2.Blo1217ORF1038* expression construct | https://benchling.com/s/seq-ZJsAzSjsy657HZ5kSEtU?m=slm-sXLa8vVT7rhNdw6VVtlK |

| | | |
|---|---|---|
| pJV465 | *MS2.Blo1217ORF1481* expression construct | https://benchling.com/s/seq-WmNO9xhyJBerBZupl4WF?m=slm-ttx2FmWCs3BfncqA23UJ |
| pJV543 | *S1.Blo1217ORF1481* expression construct | https://benchling.com/s/seq-OCPQx3imuI6k5SKnx635?m=slm-o7Ieh9KDFVLwUXJvR5Xg |
| pJV420_U | Bifidobacteria-*E. coli* plasmid with upstream barcode | https://benchling.com/s/seq-MzUfYaewZ3e0LpOdZaDH?m=slm-TnCP6CBCC9J0YMaPhe8u |
| pJV420_D | Bifidobacteria-*E. coli* plasmid with downstream barcode | https://benchling.com/s/seq-Bh6YanVKjGlXTz8W3G5a?m=slm-s2bAbKefEtCGjwnYj04E |
| pTL002 | *B. breve* RBS library parent plasmid | https://benchling.com/s/seq-vxbS915gIHMiIe3AsQI1?m=slm-I9Q0u968dmeXyQs5I9GO |

**Supplementary Table S6:** Important ssDNA used in this study.

| Primer | Sequence |
|---|---|
| linear_MTase_F | TGAGCTAACACCGTGCGTGTTGACAA |
| linear_MTase_R | CTTTCGGCGGGCTTTGCTCGAGTTA |
| pJV420_U_F | NNNNAGGCATGCAAGCTTGGCGTA |
| pJV420_U_R | GCAGGTCGACTCTAGAGGAT |
| pJV420_D_F | GGCGTAATCATGGTCATAGC |
| pJV420_D_R | NNNNAAGCTTGCATGCCTGCAG |
| HT_sequence_F | GGCTTGTGCCACAACGGTGT |
| HT_sequence_R | GGCAGTGAGCGCAACGCAAT |
| Random_RBS | GTATTTTACTGCTGGGGATTTTTATGCCCTT TGGGGCTGTANNNNNNNNNTGAAAATAATC AATATTGGAATTCTTGCCCATGTAGACGCT GGAAAGACGACCTTGACGGAGAGCCTGCT ATATG |
| RBS_screen_F | ACATATCCCCAAAAACAGCA |
| RBS_screen_R | GAGCGTCGAAAAAGGGACAA |
